# ZFP982 confers mouse embryonic stem cell characteristics by regulating expression of *Nanog, Zfp42* and *Dppa3*

**DOI:** 10.1101/2020.06.03.131847

**Authors:** Fariba Dehghanian, Patrick Piero Bovio, Zohreh Hojati, Tanja Vogel

## Abstract

We here used multi-omics analyses to identify and characterize zinc finger protein 982 (*Zfp982)* that confers stemness characteristics by regulating expression of *Nanog, Zfp42* and *Dppa3* in mouse embryonic stem cells (mESC). Network-based expression analyses comparing the transcriptional profiles of mESC and differentiated cells revealed high expression of *Zfp982* in stem cells. Moreover, *Zfp982* showed transcriptional overlap with *Yap1*, the major co-activator of the Hippo pathway. Quantitative proteomics and co-immunoprecipitation revealed interaction of ZFP982 with YAP1. ZFP982 used a GCAGAGKC motif to bind to chromatin, for example near the stemness conferring genes *Nanog, Zfp42* and *Dppa3* as shown by ChIP-seq. Loss-of-function experiments in mESC established that expression of *Zfp982* is necessary to maintain stem cell characteristics. *Zfp982* expression decreased with progressive differentiation, and knockdown of *Zfp982* resulted in neural differentiation of mESC. ZFP982 localized to the nucleus in mESC and translocated to the cytoplasm upon neuronal differentiation. Similarly, YAP1 localized to the cytoplasm upon differentiation, but in mESC YAP1 was present in the nucleus and cytoplasm.

**Graphical Abstract:** ZFP982 is a regulator of stemness of mouse embryonic stem cells and acts as transcription factor by activating expression of stem cell genes including *Nanog, Dppa3 and Zfp42*.

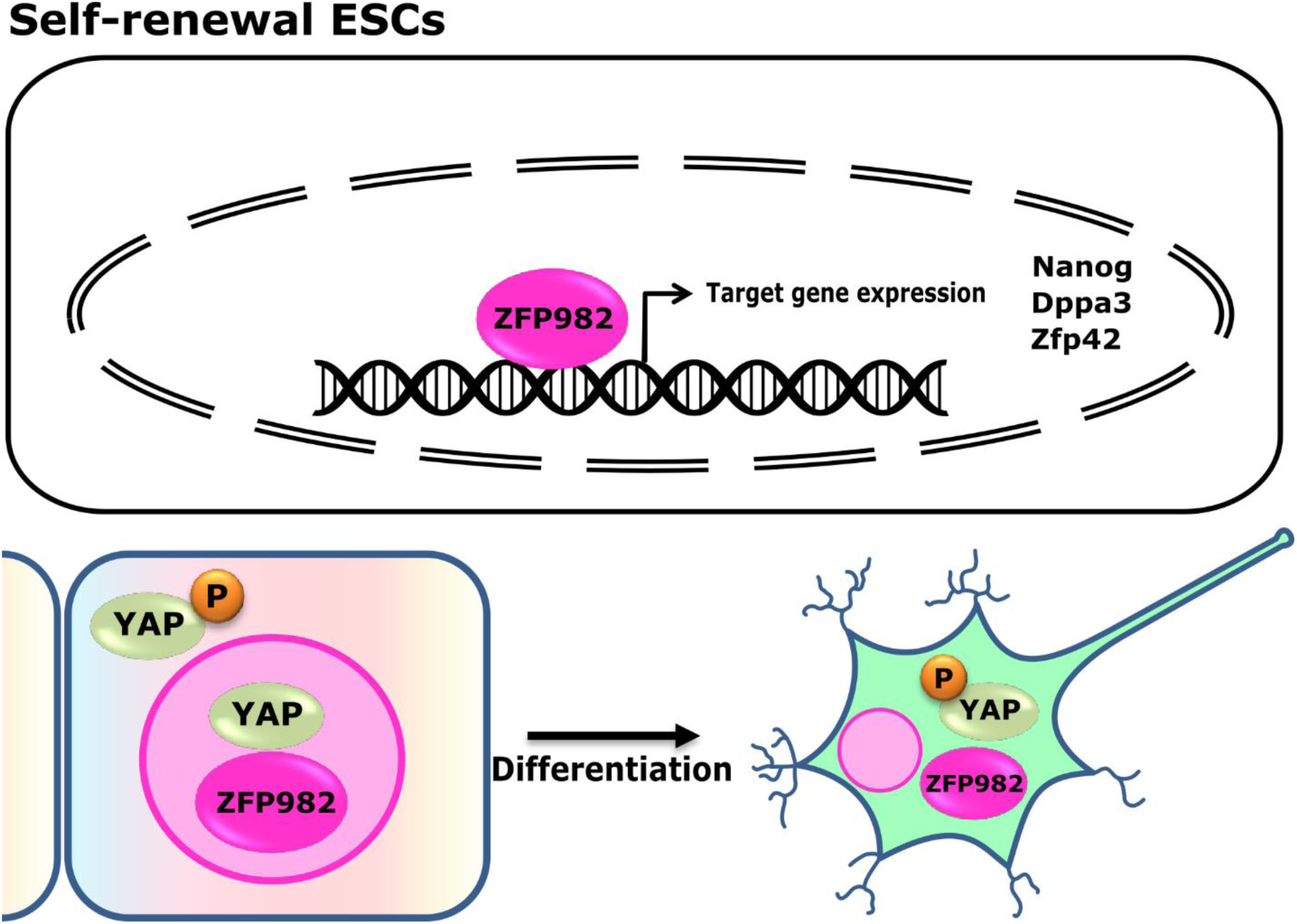

**Highlights:**

- *Zfp982* is a new mouse stem cell defining marker gene.
- *Zfp982* is co-expressed with *Yap1* and stem cell marker genes in mESC.
- ZFP982 binds to DNA and induces expression of master genes of stemness in mESC.
- Expression of *Zfp982* gene prevents neural differentiation and maintains stem cell characteristics.
- ZFP982 and YAP1 interact in mESC and translocate to the cytoplasm upon neural differentiation.

## Introduction

Embryonic stem cells (ESC) are important resources for cellular therapies, tissue repair and drug development due to their self-renewal capacity and ability to differentiate into all cell lineages [1]. ESCs derive from the inner cell mass (ICM) of the mammalian blastocyst and can be maintained *in vitro* under well-established culture conditions. Expression of stem cell specific genes and activity of various signaling pathways are important for stem cell maintenance [2]. Despite intensive research in the past, new potential components maintaining the stem cell state or inducing differentiation, respectively, are still emerging, for example by analysis of various transcriptomes. Characterizing the role of potential new genes involved in stem cell specification is critical to further advance the understanding of unprecedented mechanisms involved in ESC pluripotent maintenance and self-renewal. Moreover, such efforts bear the potential to improve potential applications of ESCs in clinical settings [3,4].

ESC fate specification is regulated by a complex transcriptional regulatory network which is controlled in response to various extrinsic signaling stimuli [5]. In recent years, a wide range of ESC markers including cell surface proteins, transcription factors, and peptides as well as specific activation of diverse signaling pathways were identified. However, still not all signaling pathways involved in ESC fate are fully explored and identification of important downstream effectors is thus of great significance [6-8]. So far, the importance of Leukemia inhibitory factor-signal transducer and activator of transcription 3 (LIF-STAT3), Transforming growth factor (TGF)-superfamily, Hippo, Wnt-β-catenin, Insulin-like growth factor (IGF) and Fibroblast growth factor (FGF) pathways has been recognized in the regulation of ESC self-renewal and pluripotency [9]. Moreover, various transcription factors are crucial for regulating the ESC transcriptome, including the zinc finger transcription factors Zinc finger protein 42 (ZFP42, REX1), Kruppel-like factor 4 (KLF4) and Sal-like protein 4 (SALL4). Combined expression of Octamer-binding transcription factor 3/4 (OCT3/4), Sex determining region Y-box 2 (SOX2), c-MYC (so-called Yamanaka factors) and these zinc finger transcription factors in somatic cells is used for reprograming the pluripotency state of stem cells [5,10]. Reprogramming of somatic cells could also be efficiently induced by the combination of OCT4, SOX2, and YAP1, the latter of which being a major co-activator of Hippo signaling pathway. YAP1 acts as a transcriptional co-activator that uses other transcription factors with the capacity of direct DNA-binding to regulate gene expression [11].

In this study, we identified the zinc finger protein ZNF982 to be highly expressed in mESC, co-expressed with stem cell markers and effectors of the Hippo pathway. ZFP982 interacted with the YAP1 protein. Chromatin immunoprecipitation followed by deep sequencing (ChIP-seq) confirmed that ZFP982 enriched in the vicinity of genes known to confer mESC stemness, including *Nanog, Zfp42* and *Dppa3*. Accordingly, we observed that increased expression levels of *Zfp982* prevented neural differentiation and maintained stem cell characteristics. In contrast, the decrease of *Zfp982* expression promoted differentiation into neural cells. Further, ZFP982 and YAP1 translocate from the nucleus to the cytoplasm upon neural differentiation. Together our data strongly promote the view that ZFP982 might serve as a transcription factor that confers mESC stemness and that associates with Hippo signaling.

## Results

### *Zfp982* is strongly transcribed in mESC and co-expressed with stem cell marker genes and members of the Hippo pathway

Maintenance of stem cell characteristics requires gene expression activities orchestrated by a variety of transcription factors. We hypothesized that other factors might be involved beyond the well-described network centered around *Oct4, Nanog* and *Sox2*.

To identify differentially expressed genes in mESC compared to differentiated cells (DC), we determined the median of the expression value for each probe set from publicly available microarray data. Each specific gene was associated with two coordinates representing median expression in the mESC data set and in the DC data set and was represented as a single dot in Figure 1A. We observed that *Zfp982*, a gene with unknown function, was highly expressed in mESC at a similar level as well-known stemness markers *Sox2, Nanog* and *Oct4* (Figure 1A).

**Figure 1:**
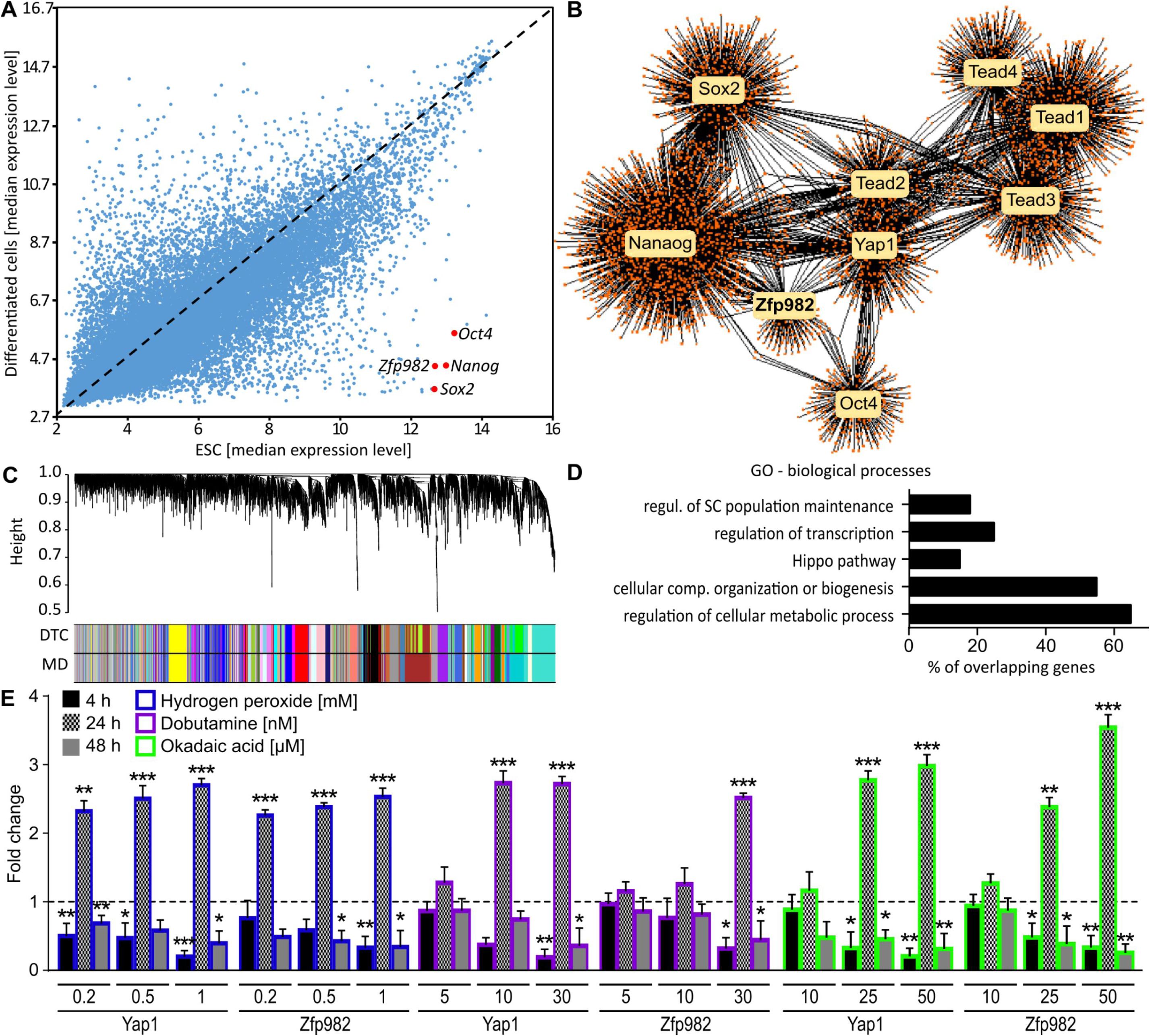
*Zfp982* gene is highly expressed in mESCs and co-expressed with stemness markers and Hippo pathway members. (A) The scatter plot of the median gene expression level in mESCs versus differentiated cells represents the high expression of *Zfp982* gene in mESCs. Each dot represents the median level of expression of each gene in mESCs (x-axis) and in differentiated cells (y-axis). Genes with higher expression level in mESCs are shown below the diagonal and genes with higher expression level in differentiated cells are detectable above the diagonal. *Oct4, Nanog* and *Sox2* genes together with *Zfp982* are highlighted. (B) Differential co-expression analysis using the Diffcorr R package indicates co-expression of *Zfp982* gene with Hippo signaling core members and stem cell markers in mESCs compared to differentiated cells. (C) Constructing a co-expression network in mESCs using WGCNA. Topological overlap-based dendrogram showing the hierarchical clustering of genes into modules. The dynamic tree-cutting (DTC) algorithm was used for module detection which is represented by a distinct color code. The presence of *Zfp982* and *Yap1* in the yellow module confirms the Diffcorr results. (D) Bar-chart of a DAVID GO term analysis for biological processes of the yellow module (p-value <0.05). (E) Hydrogen peroxide, Dobutamine and Okadaic acid induced expression changes the expression pattern of *Yap1* and Z*fp982* in P19 cells as revealed by qPCR. Significant and similar expression patterns of *Yap1* and *Zfp982* genes in response to Hydrogen peroxide, Dobutamine, and Okadaic acid were detected. For each experiment, the expression levels were normalized to control cells (untreated), which are displayed as dashed line. Mean with SEM, *: p>0.05, **: p<0.01, ***: p<0.001, unpaired Student ‘s t-test n=3.

To reveal a potential functional role of *Zfp982* in stem cells we performed co-expression and differential co-expression analyses using the DiffCorr software package, which identified differential expression correlations (0.5> (r1-r2) < - 0.5, FDR<0.05) between mESC and DC. These analyses also correlated *Zfp982* expression to mESC markers (*Oct4, Sox2, Nanog*), and additionally to core members of the Hippo pathway, *Yap1* as well as *Tead1-4* (Figure 1B), specifically in mESCs.

We next performed module identification of the co-expression network analyses identifying gene groups with similar expression patterns. Topological overlap-based dendrogram showed the hierarchical clustering of genes into modules. The dynamic tree cutting (DTC) algorithm was used for module detection (MD), which is represented by a distinct color code. Presence of *Zfp982* and *Yap1* in the yellow module confirmed the Diffcorr results (Figure 1C) of co-expression of *Zfp982* with *Yap1* together with stem cell markers in mESC but not in DC. A GO term analysis of genes represented within the yellow module indicated that they correlated with stem cell maintenance, Hippo signaling, regulation of gene expression and regulation of macromolecule biosynthetic process (Figure 1D).

Co-expression analyses correlated strongly the expression of *Zfp982* and activity of the Hippo pathway. We induced Hippo signaling in P19 cells at different time points using different concentrations of hydrogen peroxide (H_2_O_2_), dobutamine and okadaic acid (OA), and analyzed the transcription levels of *Yap1* and *Zfp982.* The treatment did not impair cell viability at these time points and by using the concentrations indicated, as we reported previously [13]. Overall, *Yap1* and *Zfp982* transcription followed similar expression patterns in response to stimulation of Hippo signaling (Figure 1E). Thereby, we observed significantly decreased expression at 4 h, significantly increased expression at 24 h, and again decreased expression after 48 h treatments for both *Yap1* and *Zfp982*. We concluded that expression of *Yap1* and *Zfp982* was most likely under control of the Hippo pathway.

### ZFP982 and YAP1 interact in mESC

To gain further insights into potential ZFP982 functions in stem cells, we performed SILAC followed by immunoprecipitation and mass spectrometric analysis of mESC overexpressing MYC-tagged ZFP982. LC–MS/MS analysis identified 161 proteins with at least two unique identified peptides, which were enriched more than 1.5 fold by MYC co-IP compared to empty vector (Figure 2A, expanded view supporting information, table 1). We analyzed the MS dataset using Cytoscape plug-in ClueGO, which revealed the most significant enriched Kyoto Encyclopedia of Genes and Genomes (KEGG)-pathway terms, for example translation initiation complex formation, metabolism of RNA, mitochondrial biogenesis and detoxification of reactive oxygen species (p-value<0.05) (Figure 2B). The involvement of ZFP982 interacting proteins in detoxification of reactive oxygen species matched our results of altered expression levels of *Zfp982* upon exposure to H_2_O_2_, dobutamine and OA.

**Figure 2.**
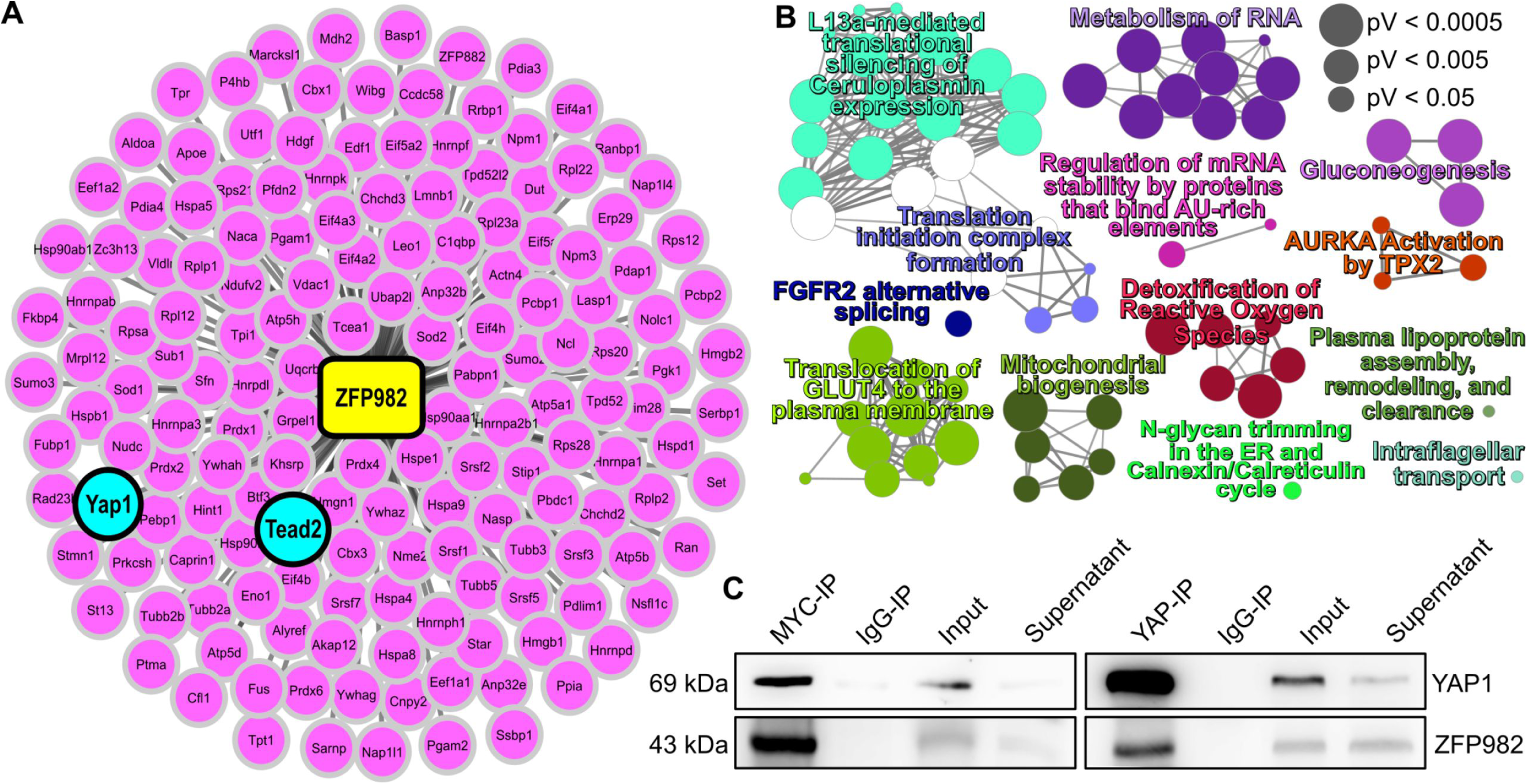
Quantitative proteomics and Co-IP reveal the interaction of ZFP982 with YAP1. (A) Protein-protein interactome of ZFP982 in mESC. Proteins with a p-value below 0.05 and fold change higher or lower than +1.5 or −1.5 were considered. (B) Gene Ontology analysis using Cytoscape (Version 3.5.1) and ClueGo plugin v2.5 for ZFP982 protein network revealed the most significant enriched Kyoto Encyclopedia of terms related to metabolism of RNA, translation initiation complex formation, detoxification of reactive oxygen species and mitochondrial biogenesis. The size of the node is related to the ClueGo generated p-value, which means that the larger nodes are more significant. (C) ZFP982 and YAP1 interact as indicated by co-IP using anti-MYC (left) or anti-YAP1 (right). IgG immunoprecipitation served as control. Input and supernatant after incubation with beads were loaded in addition. Immunoblots were probed with YAP1 (top) and ZFP982 (bottom).

Within the group of significantly enriched proteins that might associate with ZFP982 we identified YAP1 and TEAD2, both with around 4.9-fold enrichment in ZFP982-MYC IP over the empty vector condition (expanded view in supporting information, table 1). As both transcriptomic and proteomic data suggested that ZFP982 might act as an effector of the Hippo signaling pathway, we validated the protein interaction of ZFP982 and YAP1. We overexpressed ZFP982-MYC in mESC, and performed co-IP, either using anti-MYC or -YAP1 antibodies, followed by immunoblotting. These experiments confirmed that ZFP982 interacted with YAP1 in mESC (Figure 2C). Together, our data suggested so far that *Zfp982* and *Yap1* were co-expressed and that both proteins might act in concert under control of Hippo signaling.

### ZFP982 binds to chromatin and regulates expression of stemness genes in mESC

To study whether ZFP982 had DNA binding properties, we performed ChIP-seq analysis in mESC transfected either with ZFP982 overexpression vector (OE) or empty vector as control. Notably, the ChIP-seq experiment identified 7,513 regions bound by ZFP982 in mESC transfected with ZFP982-OE vector compared to the mESC transfected with empty vector (Figure 3A). From the motif analysis of ChIP-seq results, we identified a highly enriched consensus sequence of eight base pairs (GCAGAGKC; E-value= 1.8e-138) as a binding motif of ZFP982 (Figure 3B). This GCAGAGKC motif was identified in 1778 regions within the genome and our ChIP-seq peaks showed enrichment in the condition of overexpression of ZFP982 compared to the control (Figure 3C). The genome-wide distributions of peaks containing the ZFP982 motif and ChIP-peaks after ZFP982 overexpression showed mostly occurrence in distal intergenic regions, followed by intronic and promoter regions (Figure 3D).

**Figure 3.**
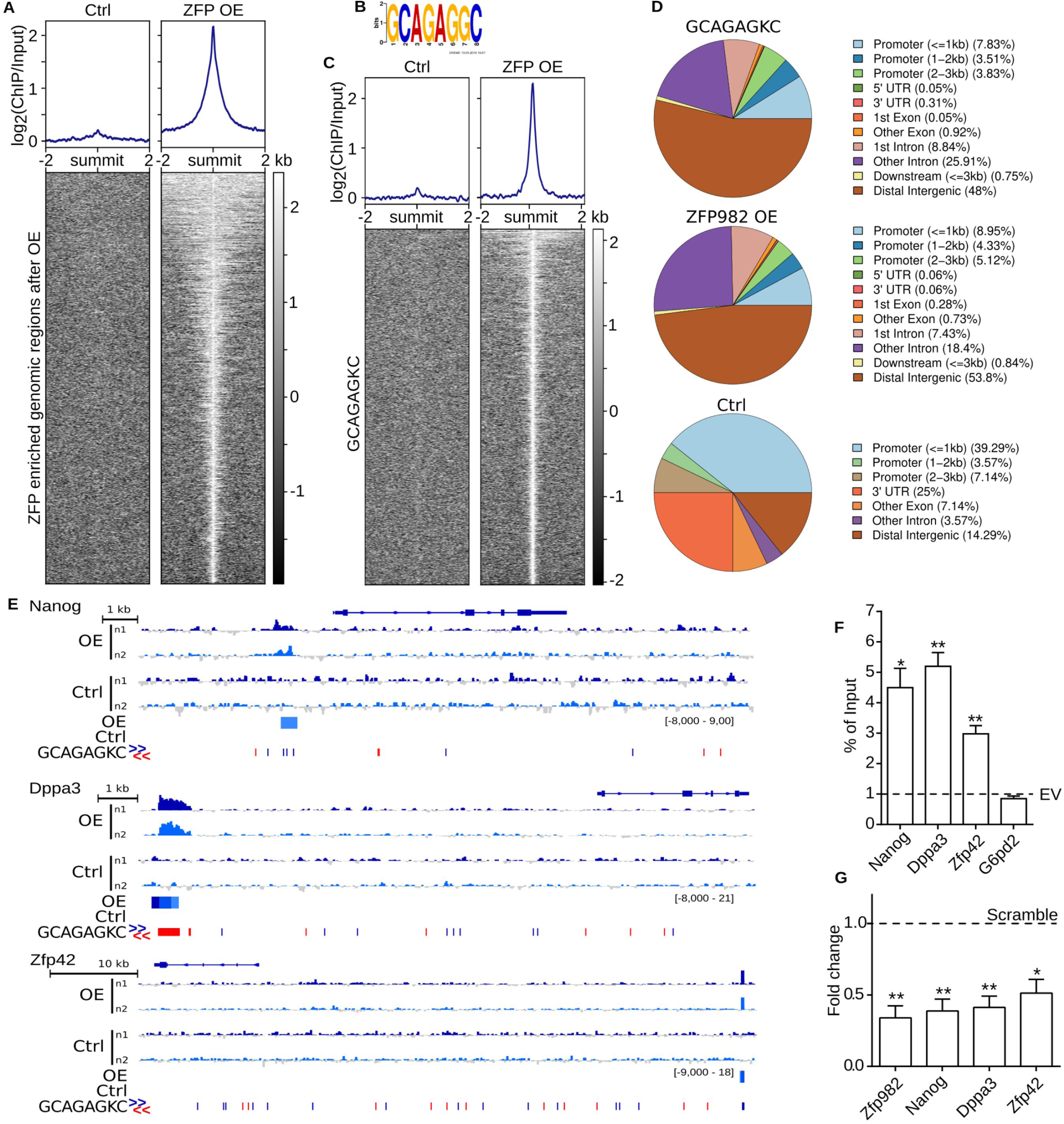
ZFP982 binds to chromatin and activates transcription of stemness genes in mESC. (A) Heatmap of ZFP982 ChIP-seq enrichment in mESCs transfected with ZFP982-OE vector and empty vector (ctrl). The heatmap shows all ZFP982 peaks ± 2 kb with each row representing a distinct peak region in OE condition. The signal intensity is displayed in SES normalized log2 (ChIP/Input). (B) Motif prediction analysis of ZFP982 peaks by MEME suite (http://meme-suite.org/). The GCAGAGKC motif is identified (E-value= 1.8e-138). (C) Peaks of ZFP982 motif GCAGAGKC. ChIP-seq heatmap of ZFP982 in mESC transfected with ZFP982-OE vector and empty vector (ctrl). The heatmap displays 1778 regions containing the motif GCAGAGKC (E-value= 1.8e-138) predicted by MEME suite (http://meme-suite.org/). Heatmap plotted ± 2 kb of the peak summit. The signal intensity is displayed in SES normalized log2 (ChIP/Input). (D) Peak annotation and visualisation performed with ChIPseeker (bioconductor chipseeker 1.14.2) showing the distribution of 1778 peaks containing the GCAGAGKC motif (top), 2130 peaks after overexpression (middle) and 48 ctrl peaks (bottom) over known genomic features in percentage. 3 kb upstream and downstream of the beginning of the peak was defined as search parameter. Flanking gene information was included within a distance of 5 kb. (E) The GCAGAGKC motif is detected in the vicinity and within *Nanog, Zfp42*, and *Dppa3* gene bodies. ChIP-seq maps of these stemness genes bound by ZFP982 in overexpression (OE) condition compared to empty vector as control (Ctrl). Two biological replicates are illustrated in each case (OE, n1-n2 and Ctrl, n1-n2) and displayed on an integrative genomics viewer (IGV, http://software.broadinstitute.org/software/igv/). Blue arrow head: binding motif in 5’ to 3’, red arrow head: motif in 3’ to 5’. (F) ChIP-qPCR to validate enrichment of ZFP982 in regulatory regions of *Nanog, Zfp42*, and *Dppa3* genes. *G6pd2* was used as negative control (no-bound region). Enrichment was displayed in the overexpression condition normalized to empty vector control (EV), indicated as dashed line. (G) *Nanog, Zfp42*, and *Dppa3* expression decreased after *Zfp982* knockdown compared to scramble control (indicated as dashed line) as detected by RT-qPCR. Data are presented as mean ± SEM. p-values were calculated using unpaired, two-tailed Student’s t-test: *p<0.05, **p<0.01, ***p<0.001, n=3.

ZFP982 was co-expressed with stem cell marker genes, i.e. *Sox2, Nanog*, and *Oct4* (Figure 1B). We therefore analyzed putative binding sites of ZFP982 in the vicinity of stemness marker genes on the chromatin. We identified several putative binding motifs within and flanking *Nanog, Zfp42* and *Dppa3*, and we detected strong enrichment of ZFP982 binding compared to the control condition in the vicinity of these stemness marker genes (Figure 3E). ChIP followed by qPCR (ChIP-qPCR) after ZFP982 overexpression in mESCs confirmed the enrichment of ZFP982 in putative regulatory regions of *Nanog, Zfp42* and *Dppa3* genes (Figure 3F).

Based on these results we examined whether ZFP982 regulated transcription of *Nanog, Zfp42* and *Dppa3*. We transfected mESCs with a scramble control, and for *Zfp982* knockdown we used the shRNA2 that showed to decrease ZFP982 levels about 60% (expanded view figure 1B). RT– qPCR showed decreased expression of *Nanog, Zfp42* and *Dppa3* genes in *Zfp982* knockdown condition compared to the control (Figure 3G). In summary, our data supported that ZFP982 might be a direct and positive regulator of self-renewal in mESC by activating transcription of pluripotency genes.

### *Zfp982* expression decreases during and prevents neural differentiation but maintains stem cell characteristics

To follow the hypothesis that ZFP982 influences stem cell characteristics and to enlighten further potential synergy with YAP1, we studied the dynamics of *Zfp982* and *Yap1* expression levels during neural differentiation. *Zfp982* expression during mESC differentiation towards neurons decreased both at protein (Figure 4A) and mRNA levels (Figure 4B). Expression of ZFP982 was hardly detectable in stages past the early radial glia like cell (RGLC) stage. *Yap1* expression decreased along differentiation as well (Figure 4A, B), but expression levels were maintained upon differentiation and detected also in the neuronal stage. The same trend of decreased expression of *Zfp982* and *Yap1* during neural differentiation was observed in P19 cells using RT-qPCR (expanded view figure 1A). In this model system of neural differentiation, *Zfp982* knockdown with shRNA2 showed that reduced expression of *Zfp982* during P19 neural differentiation significantly increased transcription of neural markers, whereas it significantly decreased expression of *Yap1*. The expression changes of the neural marker genes were comparable either after *Zfp982* knockdown (ZFP KD) or upon a differentiating retinoic acid (RA) stimulus. The combination of ZFP982 KD and RA-treatment (ZFP KD+RA) increased expression levels of neural markers and decreased expression levels of *Zfp982* and *Yap1* compared to ZFP KD or RA treatment alone (expanded view figure 1C).

**Figure 4.**
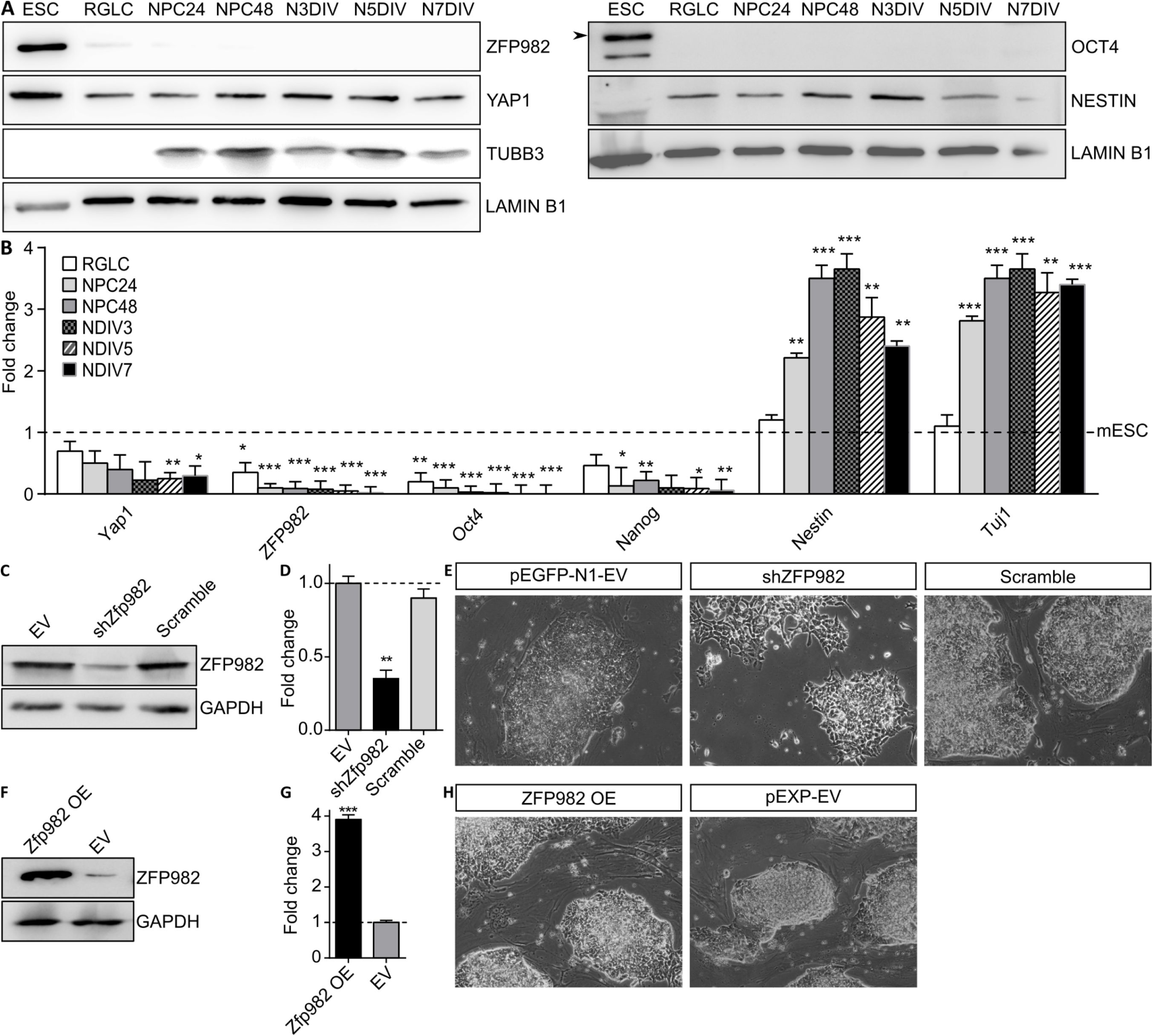
*Zfp982* expression declines during differentiation and maintains stem cell characteristics in mESCs. ZFP982 and YAP1 expression levels at different stages of V6.5 mESC differentiation assessed by immunoblot (A) and RT-qPCR (B). A significant reduction was observed for the protein expression of ZFP982. The expression level of Nestin and TUJ1 as well-known neural markers were significantly upregulated. RT-qPCR analysis of expression levels of *Zfp982, Yap1, Oct4*, and *Nanog* as stem cell markers during neuronal differentiation, as indicated by increased expression of *Nes* and *Tuj1*. Data were normalized to the mESC stage. Immunoblot (C) and RT-qPCR (D) analyses of *Zfp982* expression after *Zfp982* knockdown (shZfp982) using shRNA compared to empty vector (EV) or scramble transfected mESCs. (E) Morphological characterization through light microscopy of mESC transfected with shZfp982 compared to EV and scramble indicate the role of *Zfp982* in mESC and prevention of differentiation. *Zfp982* knockdown in mESCs leads to differentiative cellular appearance. Immunoblot (F) and RT-qPCR (G) analysis of *Zfp982* expression in ZFP982 overexpressing mESC compared to empty vector control transfections. (H) Overexpression of *Zfp982* does not alter colony morphology of mESCs compared to empty vector (pEGFP-N1-EV) condition. Mean with SEM, **: p<0.01, ***: p<0.001, unpaired Student ‘s t-test n=3. Radial glia-like cells (RGLCs), Neural Progenitor Cells after 24h (NPC24), Neural Progenitor Cells after 48h (NPC48), Neurons after 3, 5, 7 days in vitro (DIV) differentiation.

Together these data indicated that ZFP982 expression hampers neural differentiation in two different model systems. We therefore investigated morphological changes upon altered expression levels of *Zfp982* in mESC. Knockdown of *Zfp982* in mESC, which was confirmed using immunoblots and RT-qPCR (Figure 4C and D), resulted in a cell morphology characteristic for differentiated cells compared to the scramble control condition (Figure 4E). Overexpression of ZFP982, confirmed both at protein and mRNA levels (Figure 4F and G), did not alter the morphology of mESC colonies compared to empty vector (EV) control condition (Figure 4H).

In summary, these experiments indicated that 1) decreased levels of *Zfp982* coincided with neural differentiation, 2) decreased levels of *Zfp982* favored differentiation of mESC and P19 cells, and 3) *Zfp982* and *Yap1* expression were regulated through differentiative signals in the same direction. We therefore concluded that ZFP982 is a novel stem cell marker.

### Translocation of ZFP982 and YAP1 to the cytoplasm upon neural differentiation

We next used immunocytochemistry to study ZFP982 localization in mESC and differentiated cells. Because of the association of ZFP982 and YAP1 we also studied potential co-occurrence of both proteins in this cell model system. ZFP982 and YAP1 localized mainly to the nucleus both in mESC cultured on gelatin-coated plate (Figure 5A) or on inactivated MEFs (Figure 5B). Notably, upon differentiation through withdrawal of LIF, ZFP982 translocated to the cytoplasm both in mESC cultured on gelatin-coated plates (Figure 5C) or on inactivated MEFs (Figure 5D). OCT4 expression was used as a positive control stem cell marker, which was exclusively localized in the cytoplasm in mESC (Figure 5E). YAP1 localized exclusively in the cytoplasm in differentiated cells (Figure 5F), but was detectable in both nucleus and cytoplasm in mESC (Figure 5G). As expected from our previous results, differentiated neural cells did not express detectable levels of ZFP982 (expanded view Figure 2A). MEF cells were also devoid of ZFP982 immunostaining signals (expanded view Figure 2B). Together this data suggested that expression and nuclear localization of ZFP982 was characteristic for stem cells, and most likely necessary to maintain expression of stemness marker genes. ZFP982 associated and followed a similar localization dynamic as YAP1.

**Figure 5.**
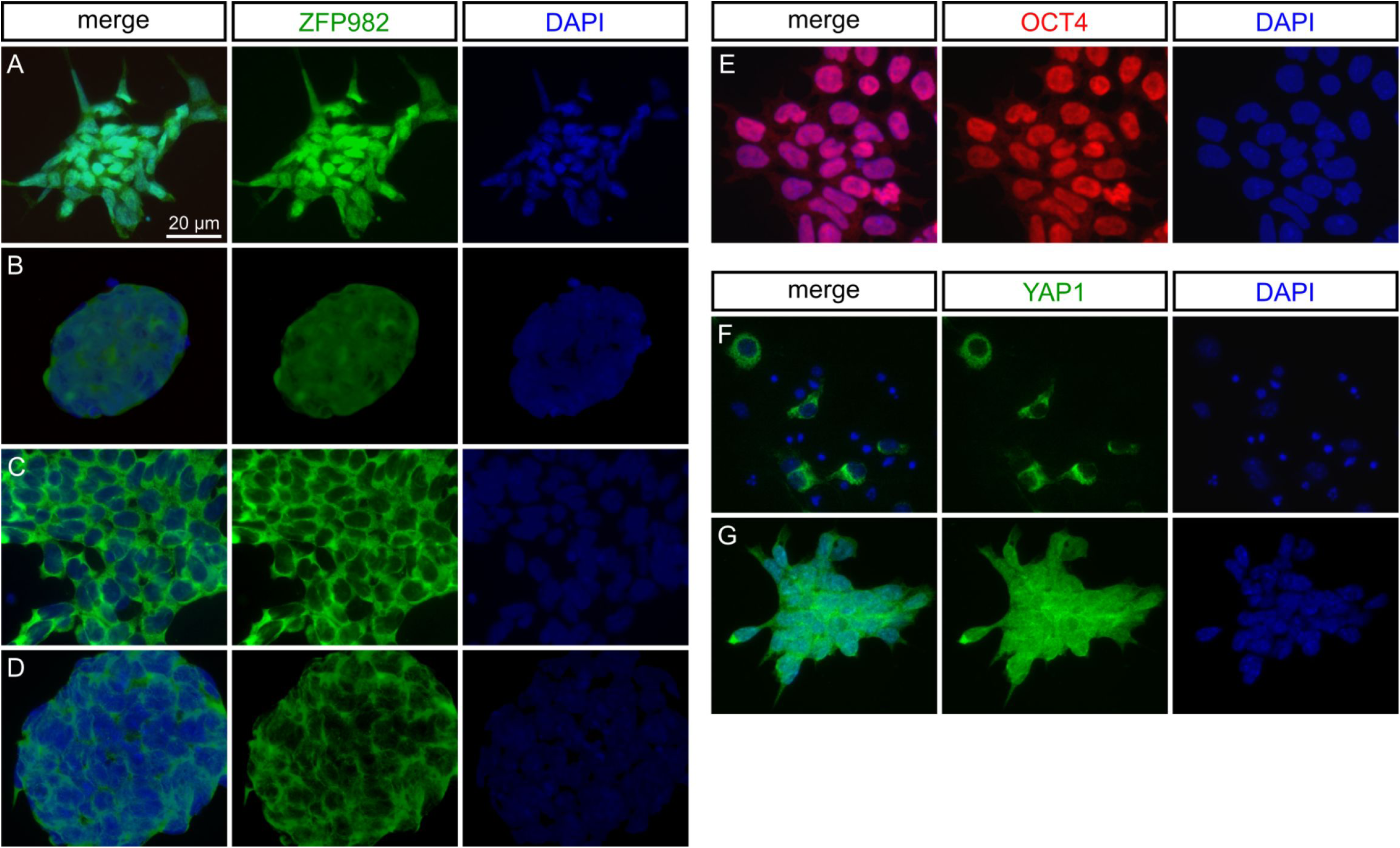
Nuclear localization of ZFP982 and YAP1 in mESC and cytoplasmic translocation upon differentiation. Nuclear localization of ZFP982 in mESCs, in both gelatin-coated (A) and iMEF cell (B) conditions in the presence of LIF. Removing the LIF factor from cell culture medium results in ZFP982 translocation to the cytoplasm in both gelatin-coated (C) and iMEF conditions (D). (E) OCT4 was used as a positive control marker of mESC. (F) YAP1 localized entirely to the cytoplasm in neural cells. (G) YAP1 is detectable both in the nucleus and cytoplasm in mESC. Leukemia inhibitory factor (LIF), Inactivated mouse embryonic fibroblasts (iMEF). Scale bar: 20μm.

## Discussion

Understanding the molecular basis of pluripotency and self-renewal in mESCs is of major interest not only for basic stem cell research but also for their usage in clinical settings. Although multiple molecular events have been identified in recent years towards better understanding the regulatory networks conferring stemness, new proteins involved in this process are still discovered. Recent research has also shown that human and mouse ESCs are different in the molecular mechanisms which regulate the pluripotency of ESCs [12,13]. mESC-specific gene expression patterns have been determined through SAGE, microarray analysis and in silico subtraction of EST database. Recently, a dataset has been prepared to investigate differentially expressed genes in mESC versus differentiated cells. In this report the authors identified E130012A19Rik (E13) as a new modulator of neural differentiation through reverse engineering analyses and experimental confirmation [14]. We used the same microarray data, which indicated that *Zfp982* was highly expressed in mESC, at similar levels of stem cell markers compared to differentiated cells. Transcriptome analyses of mESCs, however, are not sufficient to define molecular function of specific genes [15]. Therefore, we used multi-omic approaches to reveal interacting proteins and chromatin-binding properties of ZFP982, and showed involvement of ZFP982 in mouse stemness regulatory network.

OCT3/4, SOX2, NANOG, KLF4 and cMYC are essential transcription factors for maintenance of pluripotency in ESCs. Their roles as stemness factors was established through loss- and gain-of-function experiments. It has been recently reported that SOX2 is necessary for OCT3/4 expression in ESCs [16]. OCT3/4 also co-operates with SOX2 to induce transcription of the target genes such as *Sox2, Oct3/4* and *Nanog* [17]. The co-operation of OCT3/4, SOX2, NANOG, KLF4 and cMYC transcription factors to regulate expression level of each other suggests that these transcription factors form a regulatory network for maintaining stemness. The regulatory roles of *Nanog, Zfp42* and *Dppa3* genes in stem cell properties were reported in previous studies [18-20]. *Zfp42* was also identified as a gene with highly-specific expression in pluripotent stem cells. *Zfp42* expression was down-regulated after RA treatment, which induced cell differentiation [21].

The results of our study indicate that *Zfp982* is part of this network of transcriptional regulators. Multiple observations support this interpretation. First, *Zfp982* shows similar expression dynamics, high in mESC and low in differentiated cells, as the classical stemness factors. Second, ChIP-seq and ChIP-qPCR analyses added ZFP982 as a transcriptional activator of *Nanog, Zfp42* and *Dppa3* genes to the transcription factor network necessary to maintain stemness and pluripotency. Third, *Zfp982* knockdown induces neural differentiation in mESC and increased levels of *Zfp982* prevent differentiation and maintain stem cell characteristics in mESC.

Moreover, our results suggest a connection of ZFP982 to the Hippo pathway via co-expression and association of ZFP982 and YAP1. The mediation of stemness by activity of Hippo signaling and the central role of its effector YAP1 is proposed by several authors [11,22-26]. However, more recently, two studies suggested contradictory roles of YAP1 in mESCs, as these authors claimed that YAP1 is dispensable for self-renewal of ESCs [27,28]. YAP1 peaks enriched at promoters of genes important for mESCs including *Nanog, Oct4*, and *Sox2* [29], and it is therefore tempting to speculate that ZFP982 might be one of the factors that facilitate recruitment of YAP1 to the chromatin. However, we have only limited evidence that ZFP982 and YAP1 cooperate at chromatin level, also because ChIP-seq data are sparse for YAP1. *Nanog*, for which we can show the presence of ZFP982, might be a common target of YAP1 and ZFP982. Notably, we discovered protein interaction of ZFP982 with YAP1, and we also observed that ZFP982 localized to the nucleus in mESC and translocated to the cytoplasm in differentiating conditions. YAP1 was present in both nucleus and cytoplasm in mESC, but this protein localized exclusively in the cytoplasm in differentiated, neural cells. It is thus tempting to speculate that nuclear-cytoplasmic shuttling of ZFP982 might be a means to regulate YAP1 during early events of the differentiation process. ZFP982 protein level decreased remarkably during differentiation, but YAP1 protein was maintained upon mESC differentiation. Excluding ZFP982/YAP1 temporarily from the nucleus and simultaneously silencing ZFP982 expression might reduce activity of stemness-conferring transcriptional programs and allow adaptation to the functions of Hippo/YAP1 signaling in differentiated cells. However, much more research on ZFP982 and YAP1 is needed to prove functional synergy.

## Materials and methods

### Datasets, construction of co-expression networks and differential co-expression analysis

Bioinformatics analyses were performed as described previously [30]. In summary, microarray datasets GSE19836, GSE32015 (collection of 171 mESC-specific gene expression profile (GEPs) and GSE10246 (180 GEPs derived from differentiated cells (DC)) were used [14]. The preprocessing steps including quality control, background correction, normalization, and summarization were performed to obtain two expression matrices for mESC and DC. The WGCNA R package was used for construction of two signed hybrid weighted co-expression networks [31]. Next, genes were grouped into modules using the topological overlap measure TOM similarity function of the package WGCNA. Differentially co-expressed genes analysis was also performed using the Diffcorr R package [32].

### SILAC, mass spectrometry and data analysis

The V6.5 mESC line was cultured on inactivated mouse embryonic fibroblasts (MEFs) for three passages in SILAC mESC medium (expanded view figure 3). The SILAC mESC medium contained DMEM for SILAC (Thermofisher; #88364), 15 % dialyzed FBS (Thermofisher; 326400044), 1 x glutamax (Thermofisher; #35050061), 1 x non-essential amino acids (Thermofisher; #11140050), 1 x penicillin/streptomycin (Thermofisher; #15640055), leukemia inhibitor factor (Sigma Aldrich; #L5158) and ß-mercaptoethanol (Thermofisher; #31350010), supplemented with either Lys0/Arg0 (light) or Lys8/Arg10 (heavy) (0.398 mM L-arginine ^13^C_6_-^15^N_4_, 0.798 mM L-lysine ^13^C_6_-^15^N_2_, Euriso-Top, Saarbrücken, Germany). mESC were transferred to gelatin-coated plates for at least two more passages until MEFs were not visible. Light and heavy labeled mESC were each transfected using Lipofectamine LTX (#15338500, ThermoFisher Scientific, USA) and a total amount of 2 ug pf pEXP-Empty and pEXP-ZFP982 (NovaPro; #767203-1) plasmids, respectively. Transfected cells were cultured on 10 cm gelatin-coated plates in labeled medium and in presence of 700 µg/ml Geneticin (Thermofisher; #10131035) for three more passages. Both light and heavy labeled mESC were lysed in RIPA buffer (150 mM NaCl, 1 % NP-40, 0.5 % Sodium deoxycholate, 0.1 % SDS, 50 mM Tris-HCl (pH 7.4), and 1 mM EDTA) supplemented with complete protease inhibitor cocktail (#4693159001, Roche, Switzerland). The pull-down assay of ZFP982-MYC tagged protein was performed through co-Immunoprecipitation (co-IP) using MYC-Tag (9B11) antibody (#2276, Cell Signaling). For MYC-co-IP, Bradford reagent (BioRad, Germany) was used for estimation of protein amounts for both conditions. Subsequently equal amounts of protein from light and heavy labelled conditions (1 mg of each condition) were mixed in a 1:1 ratio. Then, the mix was precleared for 1h with Protein G Dynabeads (#10004D, ThermoScientific), before MYC-co-IP was done with 70 µl of MYC-coupled s Dynabeads overnight. The MYC-coupled beads were washed three times in co-IP buffer (100 mM NaCl, 20 mM Tris, 1 mM EDTA, 0.5 % NP40-alternative, pH 7.4). MYC-IP were pooled after the last washing and it was resuspended in 50 µl 1x Laemmli buffer and separated by SDS-PAGE (Supplementary Figure3). The gel was stained using coomassie and cut in 10 slices. All steps for peptide extraction were done based on the protocol reported by Rappsilber et al. 2007 [33]. LC/MS analysis was performed on an LTQ Orbitrap XL (Thermo Fisher Scientific, Bremen, Germany) either coupled to an Agilent 1200 or an Eksigent 2D nanoflow-HPLC equipped with in-house packed C18 columns of approximately 20 cm length (Reprosil-Pur 120 C-18-AQ, 3 µm (Dr. Maisch, Ammerbuch, Germany)) without a pre-column. MaxQuant (v. 1.5.3.30 (Cox and Mann 2008)) was used for analyzing raw MS data against the UniProt protein *mus musculus* sequence database (release 07.11.2013, 51,193 protein entries). For both protein lists and peptide-spectrum-matches (on modified peptides separately), a false discovery rate of 1 % was applied. Analysis of protein groups was performed with the Perseus (Tyanova et al. 2016) software. The results displayed as a ZFP982-Overexpression vector to empty vector ratio. Proteins with a p-value below 0.05 and fold change higher than +1.5 were considered. Functional enrichment analysis was performed using a Cytoscape plug-in, ClueGO. All raw data files have been deposited to the ProteomeXchange Consortium (http://proteomecentral.proteomexchange.org) with the data set identifier PXD017820.

### ChIP and ChIP-seq

Chromatin-immunoprecipitation (ChIP of ZFP982 was performed from chromatin of mESC transfected with pEXP-Empty and pEXP-ZFP982, using the MYC-Tag according to NEXSON protocol [34]. For MYC-Tag ChIP, cells were fixed for 5 min at room temperature with 1 % methanol-free formaldehyde (Thermo Scientific, #28906). Cells were lysed using ice-cold Farnham lab buffer supplemented with complete protease inhibitor cocktail (#4693159001, Roche, Switzerland). Chromatin was sheared with Bioruptor (Diagenode, Belgium) at low power for three cycles 15s on, 30 s off. For chromatin-immunoprecipitation (ChIP)-Sequencing (ChIP-seq), sequencing was performed on an Illumina HiSeq 2500, running HiSeq Control Software version 2.2.58 (single end, read length of 50 bp, 50 Mio/reads per sample). Datasets are available at gene expression omnibus (GEO) under the GSE141505 accession number. Quality control of sequencing reads was performed using FastQC [35]. CG bias and ChIP to input efficiency was controled by computeCGBias and bamFingerprint respectively [36]. Reads were then mapped to the UCSC-main RefSeq GRCm38/mm10 reference genome using Bowtie2 [37,38]. Subsequently samples were deduplicated using MarkDuplicates (http://broadinstitute.github.io/picard/). Next, signal extraction scaling (SES) was used for data normalization [39]and log2 (ratio of the number of reads) of ChIP to input was calculated by bamCompare using pseudocount of 1 and a bin size of 25 bp. ComputeMatrix, plotHeatmap and deepTools2 were used for plotting. Motif analysis was performed using MEME-ChIP version 4.11.2 [40]. The Galaxy Platform (https://usegalaxy.eu) was used for ChIP-seq analyses [41]. Further information upon standard methods (mESC culture and neural differentiation, Knockdown and overexpression of *Zfp982* gene in mESC and P19, Immunoprecipitation, Design and cloning strategies for constructing *Zfp982* shRNA, P19 culture and neural differentiation, RNA isolation, reverse transcription, and quantitative real-time PCR (qRT-PCR), Immunoblotting, Immunocytochemistry) are found in expanded view of supplementary information.

### Statistical analysis

Each experiment was performed at least three times. Statistical analysis was carried out using GraphPad Prism 6.0 software. Differences between groups were considered significant at a P< 0.05.

### Data availability

The datasets produced in this study are available in the following databases:

- Chip-Seq data: Gene Expression Omnibus GSE141505 (https://www.ncbi.nlm.nih.gov/geo/query/acc.cgi?acc=GSE141505)
- The mass spectrometry proteomics data have been deposited to the ProteomeXchange Consortium via the PRIDE [46] partner repository with the dataset identifier PXD017820.

## Acknowledgements

This research was performed at the Department of Biology, Faculty of Sciences, University of Isfahan, Isfahan, Iran. The authors would like to appreciate the financial support by the grant received from the Research Vice Chancellor of the University of Isfahan.

Bioinformatics analyses were possible through the efforts of the Freiburg Galaxy Server team.

## Author contributions

FD: designed project, performed experiments, data analysis and interpretation, manuscript writing and figure preparation; PB: performed data analyses and figure preparation; TV: conception and design of experiments, financial support, collection and assembly of data, manuscript writing; ZH: supervision of FD.

## Figure legends

**Expanded view figure 1.**
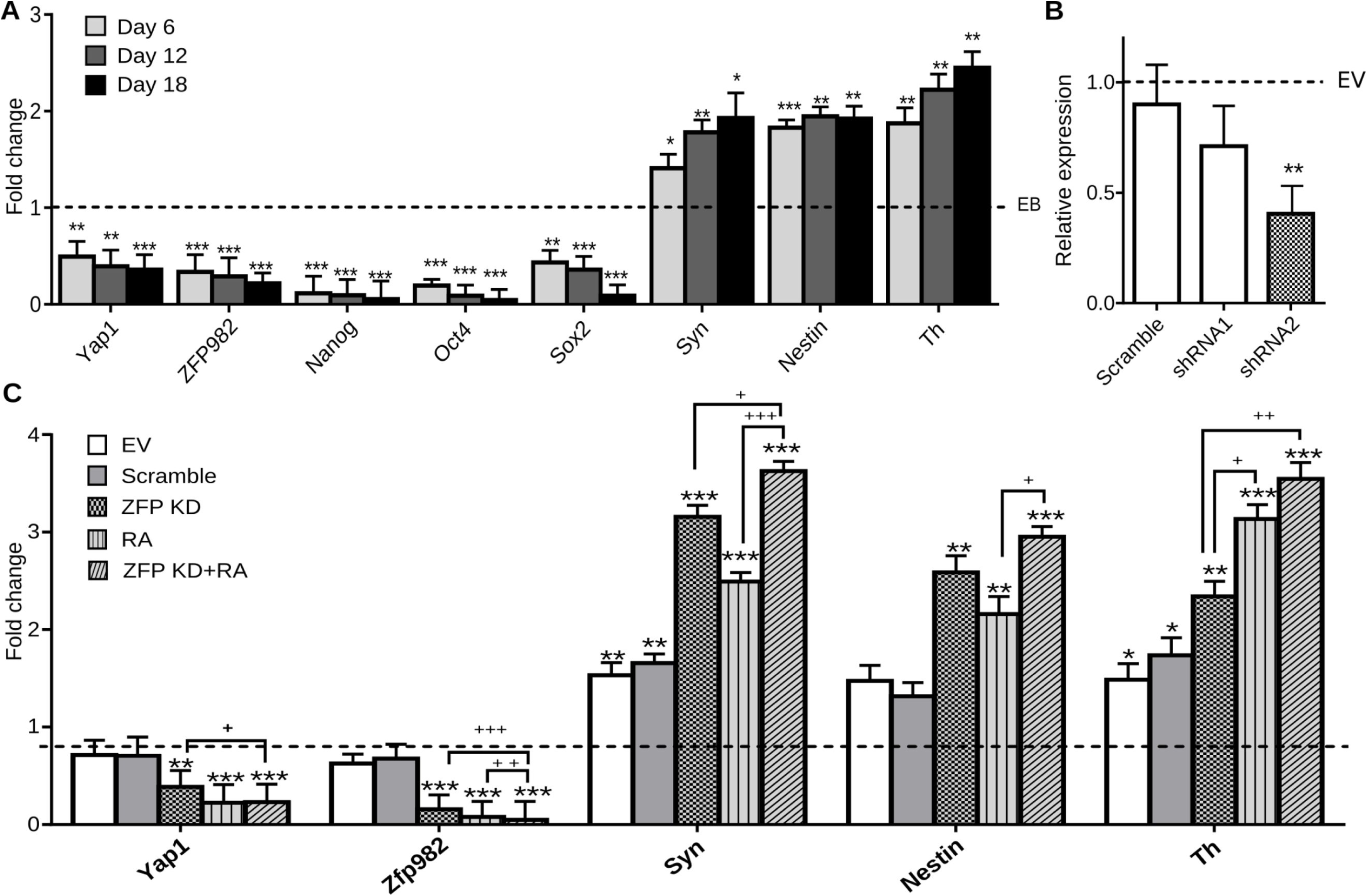
*Zfp982* expression decreases during differentiation and knockdown of *Zfp982* expression allows for neuronal differentiation in P19 cells. (A) shRNA2 significantly reduced *Zfp982* expression in transfected P19 cells compared to empty vector (EV) and Scramble control. (B) The expression level of *Zfp982* and *Yap1* gene was significantly decreased during P19 neural differentiation assessed at day 6, 12, and 18. RT-qPCR showing significant reduction of *Oct4, Sox2*, and *Nanog* expression, as well as significant up-regulation of *Nestin, Th* and *Syn* neural markers together with *Yap1* and *Zfp982* expression dynamics, which is similar to the established stem cell markers. (C) Reduced levels of ZFP982 expression during P19 neuronal differentiation leads to significantly increased expression levels of neural markers comparable to expression levels reached upon RA-treatment condition. Given is the fold change of expression of indicated marker genes as revealed by RT-qPCR in different cell stages (embryoid body (EB), retinoic acid (RA), knockdown (KD), knockdown together with retinoic acid treatment (KD+RA)). The expression levels in (B) and (C) were normalized to EB cells. Mean with SEM, * and ^+^: p>0.05, ** and ^++^: p<0.01, ***: p<0.001, unpaired Student ‘s t-test n=3.

**Expanded view figure 2:**
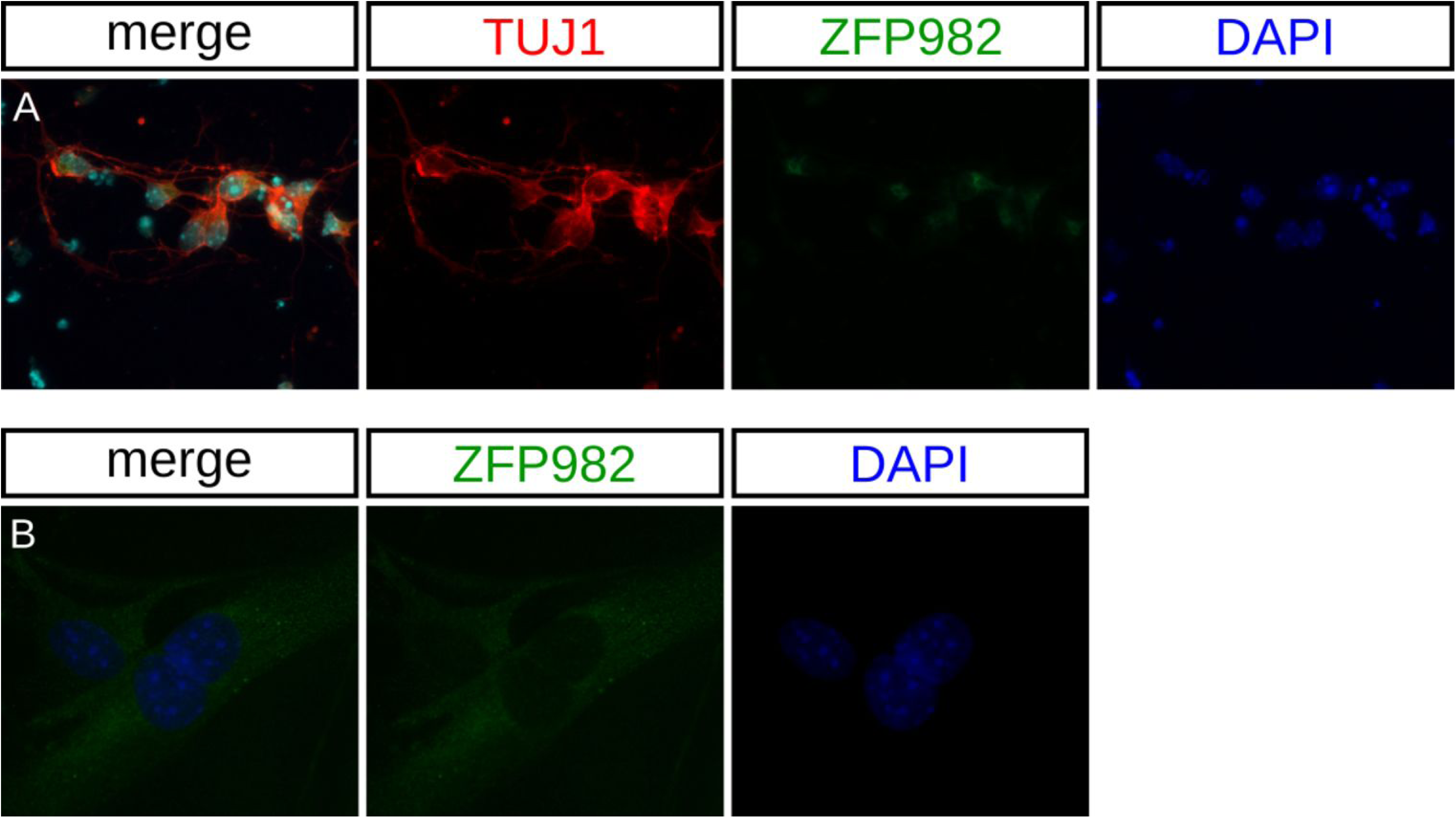
Differentiated neural and iMEF cells did not express detectable levels of ZFP982. Neural cells (A) and iMEF (B) cells which do not express ZFP982 indicate the specificity of the antibody used in mESC.

**Expanded view figure 3:**
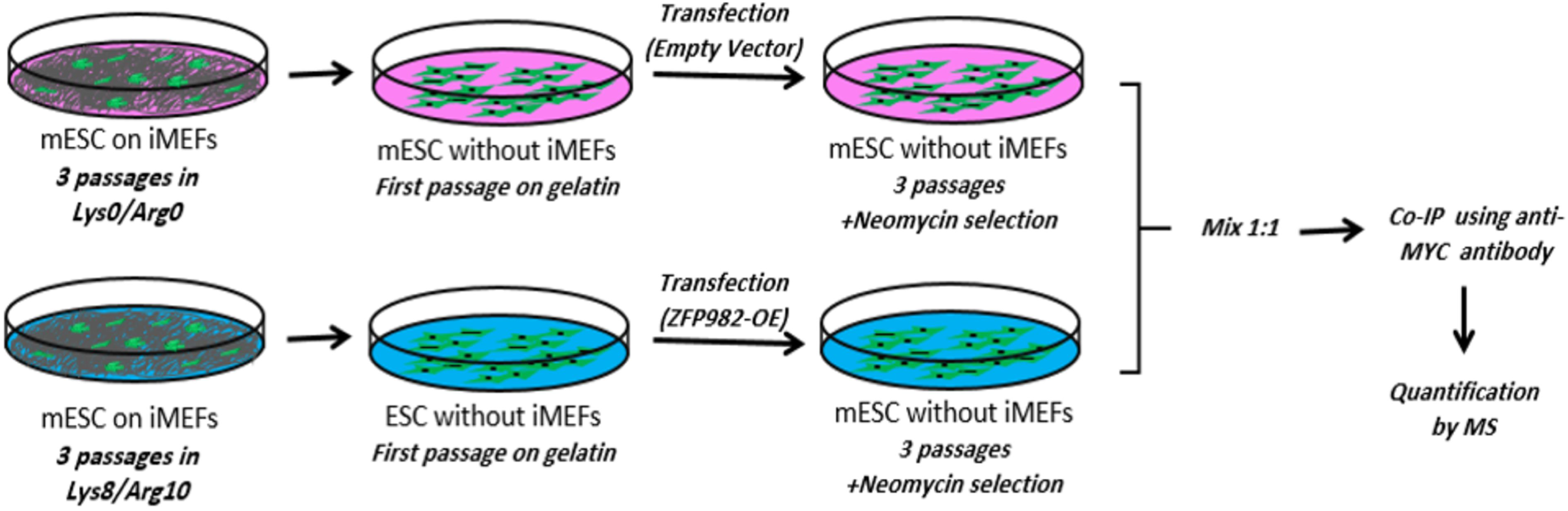
Schematic overview of mESC culture and transfection for SILAC experiment.

## References

1. Pauklin S, Pedersen RA, Vallier L (2011) Mouse pluripotent stem cells at a glance. J Cell Sci 124: 3727–3732

2. Kashyap V, Rezende NC, Scotland KB, Shaffer SM, Persson JL, Gudas LJ, Mongan NP (2009) Regulation of stem cell pluripotency and differentiation involves a mutual regulatory circuit of the NANOG, OCT4, and SOX2 pluripotency transcription factors with polycomb repressive complexes and stem cell microRNAs. Stem cells and development 18: 1093–1108

3. Bacakova L, Zarubova J, Travnickova M, Musilkova J, Pajorova J, Slepicka P, Kasalkova NS, Svorcik V, Kolska Z, Motarjemi H (2018) Stem cells: their source, potency and use in regenerative therapies with focus on adipose-derived stem cells–a review. Biotechnology advances,

4. Davidson KC, Mason EA, Pera MF (2015) The pluripotent state in mouse and human. Development 142: 3090–3099

5. Huang G, Ye S, Zhou X, Liu D, Ying Q-L (2015) Molecular basis of embryonic stem cell self-renewal: from signaling pathways to pluripotency network. Cellular and molecular life sciences 72: 1741–1757

6. Boiani M, Schöler HR (2005) Developmental cell biology: Regulatory networks in embryo-derived pluripotent stem cells. Nature reviews Molecular cell biology 6: 872

7. Hackett JA, Surani MA (2014) Regulatory principles of pluripotency: from the ground state up. Cell stem cell 15: 416–430

8. Torres-Padilla M-E, Chambers I (2014) Transcription factor heterogeneity in pluripotent stem cells: a stochastic advantage. Development 141: 2173–2181

9. Chen X, Ye S, Ying Q-L (2015) Stem cell maintenance by manipulating signaling pathways: past, current and future. BMB reports 48: 668

10. Heasley LE, Petersen BE (2004) Signalling in stem cells: meeting on signal transduction determining the fate of stem cells. EMBO reports 5: 241–244

11. Zhao Y, Fei X, Guo J, Zou G, Pan W, Zhang J, Huang Y, Liu T, Cheng W (2017) Induction of reprogramming of human amniotic epithelial cells into iPS cells by overexpression of Yap, Oct4, and Sox2 through the activation of the Hippo-Yap pathway. Experimental and therapeutic medicine 14: 199–206

12. Morey L, Santanach A, Di Croce L (2015) Pluripotency and epigenetic factors in mouse embryonic stem cell fate regulation. Molecular and cellular biology 35: 2716–2728

13. Zhao W, Ji X, Zhang F, Li L, Ma L (2012) Embryonic stem cell markers. Molecules 17: 6196–6236

14. De Cegli R, Iacobacci S, Flore G, Gambardella G, Mao L, Cutillo L, Lauria M, Klose J, Illingworth E, Banfi S (2012) Reverse engineering a mouse embryonic stem cell-specific transcriptional network reveals a new modulator of neuronal differentiation. Nucleic acids research 41: 711–726

15. Masui S, Ohtsuka S, Yagi R, Takahashi K, Ko MS, Niwa H (2008) Rex1/Zfp42 is dispensable for pluripotency in mouse ES cells. BMC developmental biology 8: 45

16. Masui S, Nakatake Y, Toyooka Y, Shimosato D, Yagi R, Takahashi K, Okochi H, Okuda A, Matoba R, Sharov AA (2007) Pluripotency governed by Sox2 via regulation of Oct3/4 expression in mouse embryonic stem cells. Nature cell biology 9: 625

17. Kuroda T, Tada M, Kubota H, Kimura H, Hatano S-y, Suemori H, Nakatsuji N, Tada T (2005) Octamer and Sox elements are required for transcriptional cis regulation of Nanog gene expression. Molecular and cellular biology 25: 2475–2485

18. Pan G, Thomson JA (2007) Nanog and transcriptional networks in embryonic stem cell pluripotency. Cell research 17: 42

19. Scotland KB, Chen S, Sylvester R, Gudas LJ (2009) Analysis of Rex1 (zfp42) function in embryonic stem cell differentiation. Developmental dynamics: an official publication of the American Association of Anatomists 238: 1863–1877

20. Bowles J, Teasdale R, James K, Koopman P (2003) Dppa3 is a marker of pluripotency and has a human homologue that is expressed in germ cell tumours. Cytogenetic and genome research 101: 261–265

21. Hosler BA, Larosa GJ, Grippo JF, Gudas L (1989) Expression of REX-1, a gene containing zinc finger motifs, is rapidly reduced by retinoic acid in F9 teratocarcinoma cells. Molecular and cellular biology 9: 5623–5629

22. Tamm C, Böwer N, Annerén C (2011) Regulation of mouse embryonic stem cell self-renewal by a Yes–YAP–TEAD2 signaling pathway downstream of LIF. J Cell Sci 124: 1136–1144

23. Bora-Singhal N, Nguyen J, Schaal C, Perumal D, Singh S, Coppola D, Chellappan S (2015) YAP1 regulates OCT4 activity and SOX2 expression to facilitate self-renewal and vascular mimicry of stem-like cells. Stem Cells 33: 1705–1718

24. Seo E, Basu-Roy U, Gunaratne PH, Coarfa C, Lim D-S, Basilico C, Mansukhani A (2013) SOX2 regulates YAP1 to maintain stemness and determine cell fate in the osteo-adipo lineage. Cell reports 3: 2075–2087

25. Lu L, Finegold MJ, Johnson RL (2018) Hippo pathway coactivators Yap and Taz are required to coordinate mammalian liver regeneration. Experimental & molecular medicine 50: e423

26. Panciera T, Azzolin L, Fujimura A, Di Biagio D, Frasson C, Bresolin S, Soligo S, Basso G, Bicciato S, Rosato A (2016) Induction of expandable tissue-specific stem/progenitor cells through transient expression of YAP/TAZ. Cell stem cell 19: 725–737

27. Chung H, Lee BK, Uprety N, Shen W, Lee J, Kim J (2016) Yap1 is dispensable for self-renewal but required for proper differentiation of mouse embryonic stem (ES) cells. EMBO reports 17: 519–529

28. Azzolin L, Panciera T, Soligo S, Enzo E, Bicciato S, Dupont S, Bresolin S, Frasson C, Basso G, Guzzardo V (2014) YAP/TAZ incorporation in the β-catenin destruction complex orchestrates the Wnt response. Cell 158: 157–170

29. Lian I, Kim J, Okazawa H, Zhao J, Zhao B, Yu J, Chinnaiyan A, Israel MA, Goldstein LS, Abujarour R (2010) The role of YAP transcription coactivator in regulating stem cell self-renewal and differentiation. Genes & development 24: 1106–1118

30. Dehghanian F, Hojati Z, Esmaeili F, Masoudi-Nejad A (2018) Network-based expression analyses and experimental validations revealed high co-expression between Yap1 and stem cell markers compared to differentiated cells. Genomics,

31. Langfelder P, Horvath S (2008) WGCNA: an R package for weighted correlation network analysis. BMC bioinformatics 9: 559

32. Fukushima A (2013) DiffCorr: an R package to analyze and visualize differential correlations in biological networks. Gene 518: 209–214

33. Rappsilber J, Mann M, Ishihama Y (2007) Protocol for micro-purification, enrichment, pre-fractionation and storage of peptides for proteomics using StageTips. Nature protocols 2: 1896

34. Arrigoni L, Richter AS, Betancourt E, Bruder K, Diehl S, Manke T, Bönisch U (2015) Standardizing chromatin research: a simple and universal method for ChIP-seq. Nucleic acids research 44: e67–e67

35. Andrews S (2010) FastQC: a quality control tool for high throughput sequence data.

36. Ramírez F, Ryan DP, Grüning B, Bhardwaj V, Kilpert F, Richter AS, Heyne S, Dündar F, Manke T (2016) deepTools2: a next generation web server for deep-sequencing data analysis. Nucleic acids research 44: W160–W165

37. Langmead B, Trapnell C, Pop M, Salzberg SL (2009) Ultrafast and memory-efficient alignment of short DNA sequences to the human genome. Genome biology 10: R25

38. Langmead B, Salzberg SL (2012) Fast gapped-read alignment with Bowtie 2. Nature methods 9: 357

39. Bailey T, Krajewski P, Ladunga I, Lefebvre C, Li Q, Liu T, Madrigal P, Taslim C, Zhang J (2013) Practical guidelines for the comprehensive analysis of ChIP-seq data. PLoS computational biology 9: e1003326

40. Machanick P, Bailey TL (2011) MEME-ChIP: motif analysis of large DNA datasets. Bioinformatics 27: 1696–1697

41. Blankenberg D, Coraor N, Von Kuster G, Taylor J, Nekrutenko A (2011) Integrating diverse databases into an unified analysis framework: a Galaxy approach. Database 2011:

