## Supplemental information for "ZFP982 confers mouse embryonic stem cell characteristics by regulating expression of *Nanog, Zfp42* and *Dppa3*"

**Supplementary information:**

**Additional Material and Methods:**

**mESC culture and neural differentiation**

Neural differentiation was performed according to the protocol of Bibel et al., 2007 [1]. Briefly, V6.5 mESC were grown on a monolayer of mytomicin-c-inactivated mouse embryonic fibroblasts (MEFs) in complete mESC medium (DMEM, 15 % FBS, 1 x non-essential amino acids, 1 x glutamax, 1 x penicillin/streptomycin, 5 μl per 500 ml ß-mercaptoethanol, 10^3^ U ml^-1^ leukemia inhibitor factor (LIF)). mESC were transferred to gelatin-coated plates after at least three passages until MEFs were not visible. To induce cell aggregation, cells grown under feeder free conditions were trypsinized and transferred to non-adherent petri dishes in complete medium for four days (DMEM, 10 % FBS, 1 x penicillin/streptomycin, 1 x non-essential amino acids, 1 x glutamax with 5 μl per 500 ml ß-mercaptoethanol). For further differentiation cells were cultivated for four days in the same medium supplemented with 5µM retinoic acid, and cell aggregates were subsequently dissociated and seeded on polyornithine/laminin coated plates in neuron precursor medium including DMEM/F12, 1 x N2 supplement, 2 mM glutamax, 1 x penicillin/streptomycin, 25µg/ml insulin, 50µg/ml BSA. The medium was replaced to a fresh medium after 2 h and one more time 24 h later. The medium was replaced by a complete neural medium (Neurobasal, 1 x glutamax, 1 x B27 supplement, 1 x penicillin/streptomycin) after 48 h of differentiation. Differentiated cells were collected at different time points including 2 h (radial glia-like cells (RGLC), 24 h (NPC24), 48 h (NPC84), 3 days (NDIV3), 5 days (NDIV5), or 7 days (NDIV7).

The following reagents for cell culture were obtained from Sigma Aldrich: polyornithine (#p3655), mytomicin C (#M0503), DMSO (#D2438), laminin (#L2020), RA (#2625), human insulin (#11376497001), BSA (#A94118), and LIF from mouse (#L5158), Trypsin (#T2600000); Thermofisher: DMEM/F12 (#11320074), DMEM (#11960044), N2 supplement (#17502048), Neurobasal (#21103049), glutamax (#35050061), non-essential amino acids (#11140050), B27 supplement (#17504044), penicillin/streptomycin (#15640055), L-glutamine (#25030081), and ß-mercaptoethanol (#31350010); Millipore: gelatin 0,1 % (#ES006B).

**Knock-down and overexpression of *Zfp982* gene in mESC and P19**

For overexpression (OE) of *Zfp982* in mESC the pEXP-Zfp982 plasmid (NovoPro, Catalog Number: 767203-1) containing the *Zfp982* cDNA was used. The pEXP-empty vector was also used as a control. Knock-down (KD in mESC and P19 cells) of *Zfp982* gene was performed using *Zfp982*-targeting shRNAs in pEGFP-N1 vector (Clontech Laboratories, Inc.). The design of shRNA1, shRNA2 and shCtrl (Scramble) in pEGFP-N1 is described in supplementary methods. For P19 transfection with sh Zfp982 and shCtrl plasmids, a total of 2 × 10^6^ P19 cells were plated in a 6 cm diameter dish 24 h before transfection. 5 μg of each expression vector were transfected by calcium phosphate precipitation [2,3]. The transfected cells were washed 16 h post-transfection and cultured in α-MEM supplemented with 10 % FBS. Following transfection, cells were treated with 500 µg/ml G418 (Invitrogen; Thermo Fisher Scientific, Inc.) for one weeks to select for successfully transfected cells expressing *Zfp982* shRNAs or scramble. RT-qPCR analysis was performed using G418-resistant clones to determine *Zfp982* mRNA expression. Neural differentiation was induced in different conditions including *Zfp982*-KD (RA)-treated, *Zfp982*-KD + RA-treated, shCtrl (Scramble) and un-transfected. For all conditions, cells were collected on day 18 of differentiation and gene expression levels, as determined by RT-qPCR, were normalized to EB stage. For KD and overexpression of *Zfp982* in mESC, 2 μg of sh Zfp982, shCtrl, pEXP-Zfp982 (ZFP982-MYC tagged expression vector) and pEXP-empty vectors were transfected using Lipofectamine 2000 (Invitrogen) according to the manufacturer's instructions. Briefly, mESC were trypsinized, pelleted by centrifugation, and resuspended into a single‐cell suspension. Subsequently, a total of 1×10^6^ mESC were combined with freshly prepared transfection complexes and plated down in 6 cm diameter dishes. Cells were treated with 700 µg/ml G418 (Invitrogen; Thermo Fisher Scientific, Inc.) for one week to select transfected cells expressing *Zfp982* shRNA, scramble, pEXP-Zfp982 and pEXP-empty vector. RT-qPCR analyses and immunoblots were performed using the G418-resistant clones to confirm overexpression and knock-down of *Zfp982* expression. The morphology of cells was also evaluated after one week.

**Immunoprecipitation**

The immunoprecipitation was performed to validate the possible interaction of ZFP982 and YAP1 proteins in mESC based on SILAC results. To confirm the interaction, the YAP1 protein was immunoprecipitated by an anti-YAP antibody and ZFP982 protein was immunoprecipitated by an anti-MYC antibody in mESC transfected with ZFP982-MYC. Transfected mESCs were lysed in co-IP buffer (100 mM NaCl, 20 mM Tris, 1 mM EDTA, 0.5% NP40-alternative, pH 7.4) supplemented with protease inhibitor. Then, the mixture was incubated for 30 min on ice, triturating every 10 min. The supernatant was collected after centrifugation (10 min, 13000 rpm). Bradford reagent (Bio-Rad) was used for determination of protein concentration. 5% input was saved and equal amounts of protein were used for both co-IPs. Protein G Dynabeads (#10004D, ThermoScientific) were coupled for 1 h at room temperature and 1 h at 4°C with Co-IP antibodies or control IgG antibody (#C15410206, rabbit IgG, Diagenode, Seraing, Belgium). Cell lysates were blocked with Protein G Dynabeads for 1 h at 4°C, subsequently transferred to antibody-coupled beads and incubated while rotating overnight at 4°C. MYC-coupled and YAP1-coupled beads were washed three times with co-IP buffer. After the last wash beads were resuspended in 30 µl 1x laemmli buffer. Finally, the complete Co-IP samples and 5% input were used for immunoblotting.

**Design and cloning strategies for constructing Zfp982 shRNA**

To create the *Zfp982* shRNA expression vector, the pEGFP-N1 vector (Clontech #6085-1) was used for DNA vector-based shRNA synthesis which was performed based on Hannon lab protocol (http://hannonlab.cshl.edu/GH_protocols.html). pEGFP-N1-Zfp982 shRNA vector expresses hairpin sequences using the H1 RNA pol III promoter, and it also has a EGFP marker under Cytomegalovirus (CMV) promoter. For each target, the sense and antisense strands were separated by a loop comprising nine nucleotides (5’TTCAAGAGA 3’) and a polythymidine tract to terminate transcription. H1 promoter sequence was also added upstream of shRNA sequence. The recognition sequence of *Ase* I restiriction enzyme was also added to the 5’ end of the H1 Promoter and 3’ end of polythymidine sequence. Two shRNA targeting *Zfp982* gene sequences were designed and synthesized. The shRNA sequences targeting Zfp982 were: shRNA1 GAAAUGCUAUACUGACAAAtt; shRNA2 TCATAAAGGAGAGAAACTTAC. The sequence of GACGCTTACCGATTCAGAA was used as the negative control scrambled shRNA (GenScript Corporation, IA) which has no significant homology to mouse or human gene. Finally, the designed sequence of 3 shRNAs were reverse complemented and sent for DNA synthesis. The oligonucleotides containing terminal *Ase* I sites were subcloned into the *Ase* I sites of the pEGFP-N1 vector to generate the pEGFP-N1 –Zfp982 shRNA vectors.

**P19 culture and neural differentiation**

The pluripotent embryonal carcinoma stem cell line P19 (Cell Bank, Pasteur Institute of Iran, Tehran, Iran) was cultured in α-minimum essential medium (α-MEM, Gibco-BRL, Carlsbad, CA, #11900073), 10 % fetal bovine serum (FBS, Gibco, #10270-106), 50 mg/ml penicillin (Sigma, #P3032) and 50 mg/ml streptomycin (Sigma, #S1277). To induce neural differentiation, cells were cultured in non-adherent plates for embryoid bodies (EBs) formation. EBs were cultured on 0.1% gelatin-coated flasks in α-MEM supplemented with 3 % FBS. The attached EBs were treated by the medium supplemented with 5×10^-7^ M retinoic acid (RA) (Sigma, #R2625). Cells were incubated at 37º C, 5 % CO_2_ and the induction medium was replaced after 24 hours. Cells were collected as EBs, on days 6, 12, 18 and utilized for further analyses and evaluation of neural differentiation.

To analyze the effects of Hippo activation we stimulated the pathway using okadaic acid (OA) (#78111-17-8, Santa Cruz, CA, USA), dobutamine (Sigma, #D0676) and Hydrogen peroxide (H_2_O_2_) (Sigma, # H1009). P19 cells were analyzed through dose- and time-response studies following exposure to different concentrations of OA (5, 10, 30 nM), dobutamine (10, 25, 50 μM) and H_2_O_2_ (0.2, 0.5, 1 mM) for 4, 24 and 48 h, respectively. Dimethyl sulfoxide (DMSO, Sigma, #D2650) was used as vehicle control for OA treatment. Normal saline was also used as vehicle control for dobutamine and H_2_O_2_.

**RNA isolation, reverse transcription, and quantitative real-time PCR (qRT-PCR)**

For RNA extraction, RNAeasy Mini Kit (#74104, Qiagen, Germany) including an on-column DNAse digestion (#79254, Qiagen, Germany) was used. The cDNA was transcribed from one µg of RNA using RevertAid MMuLV (#EP0441 Thermo Scientific, USA). The CFX-Connect Real-Time PCR detection machine (Bio-Rad, München, Germany) was used for quantitative real-time PCR analysis using GoTaq qPCR Master Mix (#A6001 Promega, Germany) based on manufacturer’s protocol. β-actin was used as a reference gene. All experiments were performed in triplicate and data analyzed by the ΔΔCt method [2]. The primer sequences were listed in the Supplementary Table S2. The GraphPad Prism 7 software and unpaired, two-tailed Student's t-test were used for statistical analyses. Data are presented as mean ± SEM.

**Immunoblotting**

Cells from different time points were harvested in Co-IP buffer (100 mM NaCl, 20 mM Tris, 1 mM EDTA, 0.5 % NP40-alternative, pH 7.4) supplemented with complete protease inhibitor cocktail (#4693159001, Roche, Switzerland). The cell lysate was incubated for 30 min on ice, triturating every 10 min 20 times. After centrifugation at 13000 rpm for 10 min, the supernatant was collected. The protein concentrations were determined using Bradford reagent (Bio-Rad). After adjustment of concentration, samples were prepared by addition of Laemmli-buffer and 5 min boiling at 95°C for SDS-PAGE. 8 % or 10 % polyacrylamide-gels were used for SDS-PAGE and proteins were transferred to PVDF membranes (Trans-blot Turbo Transfer Pack, BioRad) using the Trans-Blot Turbo (BioRad). After washing membranes in TBST (TBS with 0.1 % Tween20), membranes were incubated in 5 % BSA/TBST for 1 h. The membranes were incubated with primary antibodies in 5 % BSA/TBST, overnight at 4°C. Membranes were washed 3 times with TBST before and after incubation with the secondary HRP (horseradish-peroxidase)-coupled antibody (BioRad). The Femto reagent (Thermo Scientific) was used for membrane detection using the LAS ImageQuant System (GE Healthcare, Little Chalfont, UK). GAPDH was used as a loading control in all experiments. The following antibodies were used: anti-ZFP982 (#ab136626, 1:1000, Abcam, UK), anti-NESTIN (#ab6142, 1:1000, Abcam, UK), anti-GAPDH (#ab8245-100, 1:3000, Abcam, UK), anti-H3 (#ab12079, 1:1000, Abcam, UK), anti-YAP1 (#NB110-58358, 1:1000, Novus Biologicals, Littleton, CO), anti-OCT4 (#ab19857, 1:3000, Abcam, UK), anti-rabbit-HRP (goat, 1:10000, 115-035-003, Dianova, Hamburg, Germany), donkey-anti-mouse-HRP (goat, 1:10000, 111-005-003, Dianova). The densitometric analyses were performed with FIJI (ImageJ). Values were at first normalized to GAPDH and subsequently treated, *Zfp982*-KD and ZFP982-overexpression conditions were normalized to respective control conditions.

**Immunocytochemistry**

5 x 10^4^ cells/ml in 6 well plates were seeded on glass coverslips. Then, cells were fixed in 4 % PFA. The permeabilization and blocking of cells were done in 10 % horse serum/0.1 % Triton-X100/PBS for 1 h. Primary antibody was applied in a blocking solution at 4°C overnight. Cells were incubated with fluorophore-coupled secondary antibody at room temperature in block solution for 1 h. After 3 times washing with PBS, cells were incubated for 1 min in the DAPI solution. Coverslips were mounted on glass slides with fluorescent mounting medium (#S3023, DAKO, Jena, Germany) after 3 more washing steps in PBS. The following first and secondary antibodies were utilized: anti-ZFP982 (#ab136626, 1:500, Abcam, UK), anti-NESTIN (#ab6142, 1:1000, Abcam, UK), anti-YAP1 (#NB110-58358, 1:1000, Novus Biologicals, Littleton, CO), anti-TUJ1 (mouse, 1:100, MMS-435P, Covance), anti-OCT4 (#ab19857, 1:3000, Abcam, UK), donkey-anti-rabbit-Alexa488 (1:500, #711-545-152, Dianova) or -Alexa594 (1:500, #711-585-152, Dianova), donkey-anti-mouse-Alexa488 (1:500, #715-545-151, Dianova) or –Alexa 594 (1:500, #715-585-151, Dianova).

**Supplementary Table 1. LC–MS/MS analysis identified 161 proteins with at least two unique identified peptides, which were enriched more than 1.5 fold by MYC co-IP compared to empty vector.**

| Protein | Fold Change | P value | Protein | Fold Change | P value | Protein | Fold Change | P value | Protein | Fold Change | P value |
| --- | --- | --- | --- | --- | --- | --- | --- | --- | --- | --- | --- |
| Vldlr | 14.96 | 5.09E-295 | **Pfdn2** | 3.6431 | 1.04E-77 | **Pcbp1** | 2.7884 | 3.77E-40 | **Basp1** | 1.917 | 9.82E-25 |
| ZFP982 | 10.5 | 1.29E-288 | **Hnrnph1** | 3.634 | 5.44E-77 | **Mrpl12** | 2.7749 | 5.68E-40 | **Cnpy2** | 1.8867 | 4.47E-24 |
| Rpsa | 7.4767 | 1.80E-267 | **Hnrnpf** | 3.6249 | 3.37E-76 | **Eef1a2** | 2.754 | 6.60E-40 | **Hnrnpa3** | 1.8819 | 1.19E-23 |
| Prdx6 | 7.1214 | 2.21E-267 | **Anp32b** | 3.5635 | 4.92E-76 | **Srsf1** | 2.7513 | 7.57E-40 | **Pcbp2** | 1.8712 | 1.32E-23 |
| Chchd2 | 7.0635 | 3.92E-257 | **Sarnp** | 3.5614 | 1.11E-74 | **Atp5d** | 2.7446 | 1.06E-39 | **Npm1** | 1.8662 | 1.67E-23 |
| Btf3 | 6.1737 | 3.53E-240 | **Hmgb1** | 3.5425 | 1.42E-70 | **Eef1a1** | 2.7389 | 1.12E-39 | **Cbx1** | 1.8639 | 5.40E-23 |
| Serbp1 | 5.7793 | 4.58E-238 | **Pgam2** | 3.54 | 3.23E-68 | **Hspa4** | 2.7204 | 2.06E-39 | **Wibg** | 1.8566 | 1.02E-22 |
| ZFP882 | 5.678 | 8.36E-224 | **Eif5a** | 3.5026 | 1.11E-66 | **Eif4b** | 2.7115 | 1.00E-38 | **Set** | 1.8495 | 1.35E-22 |
| Sumo3 | 5.534 | 2.28E-211 | **Eif5a2** | 3.5026 | 1.35E-65 | **Ccdc58** | 2.6924 | 1.32E-38 | **Hnrnpd** | 1.8238 | 2.68E-22 |
| Sumo2 | 5.4733 | 1.82E-203 | **Pgam1** | 3.478 | 3.13E-64 | **Apoe** | 2.6787 | 1.70E-38 | **Leo1** | 1.791 | 6.90E-22 |
| Hsp90ab1 | 5.4085 | 1.27E-188 | **Ywhag** | 3.4765 | 3.70E-63 | **Sod1** | 2.6519 | 1.91E-38 | **Npm3** | 1.7779 | 7.10E-22 |
| Fubp1 | 5.3544 | 1.75E-185 | **Cbx3** | 3.4373 | 8.83E-63 | **Grpel1** | 2.627 | 7.31E-38 | **Utf1** | 1.7652 | 1.06E-21 |
| Nap1l1 | 5.2455 | 8.63E-176 | **Vdac1** | 3.395 | 3.79E-61 | **Caprin1** | 2.5953 | 5.71E-37 | **Star** | 1.7587 | 2.21E-21 |
| Hsp90aa1 | 5.0158 | 2.81E-175 | **Rps12** | 3.3828 | 2.53E-58 | **Tubb2a** | 2.5849 | 2.23E-36 | **Tpr** | 1.7565 | 2.61E-21 |
| Dut | 4.9821 | 3.00E-174 | **Hspa8** | 3.3761 | 2.70E-57 | **Tubb2b** | 2.5849 | 2.99E-36 | **Nolc1** | 1.7224 | 2.67E-21 |
| Pdap1 | 4.927 | 7.12E-170 | **Pgk1** | 3.3244 | 1.44E-56 | **Tubb3** | 2.5849 | 7.67E-36 | **Alyref** | 1.6519 | 4.04E-21 |
| Yap1 | 4.91 | 2.62E-166 | **Srsf2** | 3.2714 | 3.67E-56 | **Tubb5** | 2.5849 | 1.54E-35 | **P4hb** | 1.6067 | 1.13E-20 |
| Tead2 | 4.89 | 3.50E-163 | **Rps20** | 3.2304 | 1.16E-55 | **Zc3h13** | 2.5709 | 6.26E-35 | **Hnrnpa2b1** | 1.5995 | 1.16E-20 |
| Ppia | 4.8683 | 3.11E-159 | **Tpi1** | 3.2186 | 2.99E-55 | **Khsrp** | 2.5404 | 1.00E-33 | **Pdia3** | 1.5937 | 1.55E-20 |
| Nasp | 4.8587 | 9.75E-158 | **Tcea1** | 3.2155 | 1.27E-52 | **Hnrnpa1** | 2.525 | 2.46E-33 | **C1qbp** | 1.5894 | 1.90E-20 |
| Sub1 | 4.7214 | 2.34E-152 | **Fus** | 3.1936 | 5.23E-51 | **Hsp90b1** | 2.432 | 2.53E-33 | **Pdlim1** | 1.5673 | 2.20E-20 |
| Ptma | 4.6261 | 3.81E-144 | **Nsfl1c** | 3.1594 | 3.58E-50 | **Pabpn1** | 2.4134 | 5.10E-32 | **Hspb1** | 1.5409 | 2.30E-20 |
| Ranbp1 | 4.6079 | 2.04E-138 | **Prdx4** | 3.1358 | 3.91E-50 | **Rad23b** | 2.3674 | 5.94E-32 | **Sod2** | 1.5269 | 2.64E-20 |
| Ywhah | 4.5905 | 1.46E-123 | **Rplp2** | 3.0923 | 6.45E-47 | **Rps28** | 2.3181 | 6.10E-32 | **Rpl22** | 1.5245 | 1.03E-19 |
| Pbdc1 | 4.5092 | 1.58E-123 | **Hdgf** | 3.0736 | 1.81E-46 | **Hnrpdl** | 2.2902 | 6.10E-32 | **Hmgn1** | 1.51 | 1.33E-19 |
| Nme2 | 4.448 | 3.03E-123 | **Lmnb1** | 3.0535 | 2.37E-46 | **Hspe1** | 2.27 | 2.95E-31 | **Hspa5** | 1.5889 | 1.58E-19 |
| Stip1 | 4.3619 | 9.69E-121 | **Pebp1** | 3.0351 | 9.32E-46 | **Lasp1** | 2.2355 | 5.55E-31 | **Ncl** | 1.5617 | 3.41E-19 |
| Naca | 4.3523 | 8.32E-119 | **Ran** | 3.0231 | 9.38E-46 | **Rps21** | 2.2326 | 1.30E-30 | **Chchd3** | 1.536 | 4.75E-19 |
| Tpt1 | 4.3466 | 8.72E-118 | **Mdh2** | 3.0212 | 2.31E-45 | **Srsf5** | 2.2264 | 2.43E-30 | **Eif4a2** | 1.53261 | 8.84E-19 |
| Hmgb2 | 4.3113 | 6.60E-106 | **Prdx1** | 3.015 | 2.42E-45 | **Rpl23a** | 2.1957 | 1.90E-29 | **Ndufv2** | 1.53251 | 3.84E-18 |
| St13 | 4.2916 | 1.82E-97 | **Sfn** | 3.0017 | 5.49E-45 | **Srsf3** | 2.1921 | 3.39E-28 | **Rrbp1** | 1.53181 | 2.64E-17 |
| Prdx2 | 4.2895 | 1.85E-96 | **Ywhaz** | 2.9879 | 3.15E-44 | **Stmn1** | 2.1741 | 4.35E-28 | **Srsf7** | 1.5025 | 7.21E-17 |
| Eif4h | 4.2425 | 7.81E-96 | **Hspd1** | 2.9703 | 8.72E-44 | **Actn4** | 2.146 | 2.36E-27 | **Eif4a1** | 1.5 | 7.44E-17 |
| Edf1 | 3.9569 | 1.20E-95 | **Hint1** | 2.931 | 1.71E-42 | **Tpd52l2** | 2.1019 | 3.76E-27 | **Eif4a3** | 1.5 | 8.84E-17 |
| Atp5b | 3.8652 | 1.75E-93 | **Atp5h** | 2.9004 | 2.85E-42 | **Akap12** | 2.0991 | 7.79E-27 | **Hnrnpab** | 1.5856 | 9.46E-17 |
| Rplp1 | 3.86 | 5.89E-92 | **Eno1** | 2.8997 | 4.42E-42 | **Hnrnpk** | 2.0908 | 1.14E-26 | **Hspa9** | 1.5575 | 1.10E-16 |
| Nudc | 3.8483 | 6.92E-85 | **Fkbp4** | 2.8681 | 1.08E-41 | **Trim28** | 2.0345 | 3.32E-26 | **Prkcsh** | 1.5551 | 1.24E-16 |
| Anp32e | 3.8234 | 1.95E-84 | **Cfl1** | 2.8447 | 3.32E-41 | **Pdia4** | 2.0108 | 2.00E-25 | **Erp29** | 1.5166 | 1.58E-16 |
| Nap1l4 | 3.8222 | 9.47E-84 | **Ubap2l** | 2.831 | 7.20E-41 | **Uqcrb** | 2.0108 | 2.29E-25 |  |  |  |
| Tpd52 | 3.7934 | 8.38E-81 | **Aldoa** | 2.815 | 1.53E-40 | **Ssbp1** | 2.0083 | 4.52E-25 |  |  |  |
| Marcksl1 | 3.698 | 5.13E-79 | **Rpl12** | 2.8031 | 3.09E-40 | **Atp5a1** | 1.9774 | 4.67E-25 |  |  |  |

**Supplementary Table 2: Chip-qPCR and qPCR Primers**

| **Chip-qPCR primers** | |
| --- | --- |
| G6pd2 Fp (negative control) | ACAGTCTATGAAGCAGTCAC |
| G6pd2 Rp (negative control) | CTCCACTATGATGCGGTTA |
| Zfp42 Fp | AACACAATGTCAAAAGCCACAC |
| Zfp42 Rp | TGTGCATCACTACACCCAG |
| Dppa3 Fp | AGCAGTAGTGGTGCATGCCTGG |
| Dppa3 Rp | GGCAGAGGCAGAGGCAGAGGCA |
| Nanog Fp | CCGCAGAGGCAGTGGTAACAA |
| Nanog Rp | ACCACTGCCACCATCGTCAC |
| **qPCR primers** | |
| Yap1-Fp | GTCCTCCTTTGAGATCCCTGA |
| Yap1-Rp | TGTTGTTGTCTGATCGTTGTGAT |
| Zfp982 Fp | TAATCCGGGCAACAAGAATG |
| Zfp982 Rp | GCACAATGACCTCTGAGCAA |
| Nestin Fp | TCCGGGCCCCTGAAGTCGAG |
| Nestin Rp | CCAGGGCTTCCACAGCCAGC |
| Syn Fp | CATTCATGCGCGCACCTCCA |
| Syn Rp | TTGCTGCCCATAGTCGCCCT |
| Th Fp | ACCTGGGAACCCATTGGAGGCT |
| Th Rp | TCTGGCACTGCGCACATCGT |
| Oct4 Fp | TTTCTGAAGTGCCCGAAGCCCT |
| Oct4 Rp | TTCCATAGCCTGGGGTGCCAAA |
| Sox2 Fp | CACATGAAGGAGCACCCGGATT |
| Sox2 Rp | ATCATGCTGTAGCTGCCGTTGC |
| Nanog Fp | TGGGAACGCCTCATCAATGCCT |
| Nanog Rp | CGCATCTTCTGCTTCCTGGCAA |
| Tuj1(Tubb3) Fp | GGTGGACTTGGAACCTGGAA |
| Tuj1(Tubb3) Rp | TAAAGTTGTCGGGCCTGAATTCAA |
| Dppa3 Fp | CCAAGAGAAGGGTCCGCACTTT |
| Dppa3 Rp | GCAGAGACATCTGAATGGCTCAC |
| Zfp42 Fp | TTGCCTCGTCTTGCTTTAGG |
| Zfp42 Rp | AAAATGAATGAACAAATGAAGAAAA |
| Gapdh Fp | CGGCCGCATCTTCTTGTG |
| Gapdh Rp | TGACCAGGCGCCCAATAC |

1. Bibel M, Richter J, Lacroix E, Barde Y-A (2007) Generation of a defined and uniform population of CNS progenitors and neurons from mouse embryonic stem cells. *Nature protocols* **2:** 1034

2. Livak KJ, Schmittgen TD (2001) Analysis of relative gene expression data using real-time quantitative PCR and the 2− ΔΔCT method. *methods* **25:** 402-408

3. Jordan M, Schallhorn A, Wurm FM (1996) Transfecting mammalian cells: optimization of critical parameters affecting calcium-phosphate precipitate formation. *Nucleic acids research* **24:** 596-601
